# Exposure to a gradient of warming and acidification highlights physiological, molecular, and skeletal tolerance thresholds in *Pocillopora acuta* recruits

**DOI:** 10.1101/2025.01.08.632024

**Authors:** Jill Ashey, Federica Scucchia, Ariana S. Huffmyer, Hollie M. Putnam, Tali Mass

**Affiliations:** University of Rhode Island, Department of Biological Sciences, Kingston, RI 02881, USA; University of Haifa, Department of Marine Biology, Leon H. Charney School of Marine Sciences, Haifa, 3498838, Israel; University of Washington, School of Aquatic and Fisheries Science, Seattle, WA 98195, USA

**Author notes:** corresponding author: Jill Ashey. equal contribution to authorship (Jill Ashey, Federica Scucchia).

**Keywords:** transcriptomics, biomineralization, thermal stress, ocean acidification, environmental fluctuations

## Abstract

Ocean warming and acidification are among the biggest threats to the persistence of coral reefs. Organismal stress tolerance thresholds are life stage specific, can vary across levels of biological organization, and also depend on natural environmental variability. Here, we exposed the early life stages of *Pocillopora acuta* in Kāne‘ohe Bay, Hawai‘i, USA, a common reef-building coral throughout the Pacific, to projected ocean warming and acidification scenarios. We measured ecological, physiological, biomineralization, and molecular responses across the critical transition from larvae to newly settled recruits following 6 days of exposure to diel fluctuations in temperature and pH in Control (26.8-27.9°C, 7.82-7.96 pH_Total_), Mid (28.4-29.5°C, 7.65-7.79 pH_Total_) and High conditions (30.2-31.5°C, 7.44-7.59 pH_Total_). We found that *P. acuta* early life stages are capable of survival, settlement, and calcification under all scenarios. The High conditions, however, caused a significant reduction in survival and settlement capacity, with changes in the skeletal fiber deposition patterns. In contrast to a limited impact on the expression of biomineralization genes, the dominant transcriptomic response to the High conditions relative to the two other treatments included depressed metabolism, reduced ATP production and increased activity of DNA damage-repair processes. Collectively, our findings indicate that corals living in environments with large diurnal fluctuations in seawater temperature and pH, such as Kāne‘ohe Bay, can tolerate exposure to moderate projected increased temperature and reduced pH. However, under more severe environmental conditions significant negative effects on coral cellular metabolism and overall organismal survival jeopardize species fitness and recruitment.

## Introduction

The environmental impacts of climate change, especially global warming and ocean acidification (OA), are threatening coral reefs worldwide (Hughes et al., 2017). Significant research effort has been devoted to understanding how these environmental stressors will impact physiological functioning of corals (Van Woesik et al., 2022) and the dynamics of the entire reef ecosystem (Voolstra et al., 2023). Rising seawater temperatures challenge the symbiotic partnership of the coral holobiont (Helgoe et al., 2024), physiological processes (Krämer et al., 2022), and fitness (Riegl & Purkis, 2015; Hughes et al., 2019). While marine heatwaves are increasingly driving rapid mass mortality of corals (Shlesinger & Van Woesik, 2023), OA can occur simultaneously and alter coral metabolism by creating an acid-base imbalance within the coral tissues (Gibbin et al., 2015; Klein et al., 2022). OA disrupts the conformation and activity of enzymes, such as carbonic anhydrases and alkaline phosphatases, involved in host-symbiont carbon exchange and calcification (Dubinsky & Stambler, 2011). In addition, under OA conditions skeleton deposition requires more energy to remove excess H⁺ from the calcifying space (Ries, 2011), thereby posing a chronic energetic strain. Such energetic strain would further be exacerbated by the loss of nutrition derived from translocated carbon products from symbiotic algae under thermal stress (Tremblay et al., 2016; Sun et al., 2020; Rädecker et al., 2021; Allen-Waller & Barott, 2023). In light of this, studies examining coral response to these multiple and interactive stressors, in energetically sensitive early life stages are paramount to predict future coral survival.

Physiological effects from multiple stressors can be more pronounced on the early life stages of stony corals, due to the high amount of energy required for actively swimming, settling, metamorphosing and initiating calcification (Harii et al., 2002, 2010; Edmunds et al., 2013). Studies that have explored the combined effects of warming and acidification on newly released larvae and recruits have found both negative responses and signs of resilience (Albright, 2011; Byrne & Przeslawski, 2013; Putnam et al., 2013). For example, larvae and recruits exposed to combined warming and OA exhibit reductions in parameters such as survival (Cumbo et al., 2013), protein content (Cumbo et al., 2013), biomass (Anlauf et al., 2011), respiration (Cumbo et al., 2013) and skeletal weight (Foster et al., 2015). In contrast, no changes in survival (Anlauf et al., 2011; Foster et al., 2015; Sun et al., 2020, 2024), settlement (Anlauf et al., 2011; Foster et al., 2015; Sun et al., 2024), size (Bahr et al., 2020), photophysiology (Putnam et al., 2013) and respiration (Putnam et al., 2013) have been observed in response to the combination of higher temperature and acidified seawater. While some of these conflicting results may be due to magnitude and duration of exposure relative to local conditions (Kenkel et al., 2013; Kenkel & Matz, 2017; Jury & Toonen, 2019; Kurihara et al., 2021), or to differences in species sensitivity (Loya et al., 2001; Fitt et al., 2009; Comeau et al., 2014; Porro et al., 2023), it is clear that we have much to learn through integrative cross-biological scale investigations of multiple warming and OA regimes to determine organismal tolerance thresholds to environmental stress.

To better understand the complexity of physiological responses, it is critical to explore underlying cellular processes, pathways and mechanisms (Pan et al., 2015), such as through the assessment of gene expression patterns (Evans & Hofmann, 2012). Transcriptomics can reveal changes in gene expression to various environmental perturbations, defining the limits of adaptation and physiological thresholds (Gracey & Cossins, 2003; Palumbi et al., 2014; DeBiasse & Kelly, 2016). Studies evaluating the gene expression underlying the response to combined warming and acidification in early coral life stages are more scarce than the adult life stage, with some studies conducted on larvae (Putnam et al., 2013; Rivest et al., 2018; Jiang et al., 2022)), and only one study focused on combined warming and acidification stress on newly settled coral recruits (Sun et al., 2020). Coral settlement is a crucial stage at which calcification is initiated, underlined by increased expression of genes of the ‘biomineralization toolkit’ (Mass et al., 2016). Successful settlement of young corals is fundamental for the overall reef resilience (Hughes & Connell, 1999; Hughes & Tanner, 2000). Therefore, the transcriptomic and skeletal consequences of initiating calcification under OA and warming conditions urgently calls for further investigation.

The synergistic effects of climate change stressors will also be determined by species-or population-specific tolerance thresholds (Staudt et al., 2013; Pascual et al., 2022; Dilworth et al., 2024), which are shaped by present-day natural environmental regimes (Ainsworth et al., 2016; Wall et al., 2021) through acclimatization and local adaptation (Van Oppen et al., 2015; Boyd et al., 2016). Mounting evidence shows that high frequency environmental fluctuations encourage greater tolerance to stress, by, for example, reducing coral bleaching incidence under warmer temperatures (Safaie et al., 2018), increasing constitutive gene expression in corals from areas of higher thermal variability (Barshis et al., 2013), improving coral ability to regulate cellular acid-base homeostasis (Brown et al., 2022), boosting growth of coral recruits under acidic conditions (Dufault et al., 2012), and overall shaping plastic phenotypic responses across marine phyla (Fusi et al., 2024). Collectively, these studies indicate that the integration of environmental fluctuations into experimental studies is critical to allow better forecasting of organism vulnerability and ecosystem resilience under climate change conditions (Rivest et al., 2017; Kroeker et al., 2020; Tanvet et al., 2023). In Kāne‘ohe Bay, Hawai‘i, the benthic feedbacks and long seawater residence time tend to drive large natural diel fluctuations in temperature and pH (Drupp et al., 2011; Jury et al., 2013). Such daily changes in temperature and pH range from 0.5-2.5°C and 0.06-0.46 units (total scale), respectively (Guadayol et al., 2014). This diurnal variability highlights the need to incorporate environmentally relevant fluctuations in the context of the coral response to future climate. Here, we used daily changes in temperature and pH as setting for interpreting the biological response of early life stages of *Pocillopora acuta*, a common reef-building coral in Kāne‘ohe Bay, to projected future ocean scenarios. We cultured newly released larvae and primary polyps (metamorphosed and settled larvae) of *P. acuta* at ambient (daily minimum to maximum; 26.8 to 27.9°C, 7.82 to 7.96 pH_Total_), Mid (28.4 to 29.5°C, 7.65 to 7.79 pH_Total_) and High conditions (30.2 to 31.5°C, 7.44 to 7.59 pH_Total_). We then employed an integrative approach across multiple biological scales including molecular (host gene expression), morphological (coral skeletal formation), and physiological/ecological (settlement and survivorship). Because larvae and newly settled polyps are critical to population connectivity and recovery, it is important to understand the degree of susceptibility to the combined effects of warming and OA in these energetically demanding early life stages, while simultaneously accounting for natural daily environmental fluctuations.

## Methods

### Adult collection and larval release

Fifteen adult *P. acuta* colonies (hermaphroditic brooder with vertical transmission of symbionts to offspring) were collected from the Hawai’i Institute of Marine Biology (HIMB) Moku o Loʻe fringing reef (Reef #1; 21°26′03″ N, 157°47′12″ W; Figure S1A) on June 11, 2022 (four days after the first quarter moon) under a Special Activity Permit (SAP 2022-28, Division of Aquatic Resources, Honolulu, HI). Colonies (10-15 cm in diameter) with at least 5 meters between them were collected from a depth of 1-2 meters using a hammer and chisel. Colonies were returned to HIMB and placed in two 510-L outdoor mesocosm tanks (8 colonies in one tank and 7 in the other tank) with ambient flow-through seawater. Tanks were covered with shade cloth and experienced a natural solar cycle, with a diel range of 0 to 240 µmol photons m^-2^ s^-1^ (Apogee Underwater Quantum Flux meter MQ-510, Apogee Instruments).

In Hawai’i, *P. acuta* typically releases larvae on lunar days around the first quarter moon and full moon (Richmond & Jokiel, 1984). *Pocillopora acuta* adult colonies were prepared to track and capture larval release on July 6-July 11 (quarter moon on July 6, full moon on July 13). In the evenings (∼19:00 HST), individual adult colonies were placed in 2.8-L bowls and ambient unfiltered seawater was provided into the bowls at flow rate of ∼100 mL min^-1^. The water outflow of each bowl was directed into 1-L tripours with 153 µm mesh bottoms. Larvae released from the adult colonies flowed out of the bowl and into the mesh-bottomed tripour, where they were collected the following morning (∼06:00 HST). On July 10th, 2022, 9 of the 15 colonies released larvae and larvae were pooled amongst all colonies and allocated to treatments.

### Experimental set up and larval exposures

Seawater temperature is predicted to increase by ∼0.2-3.5 °C with concurrent reductions in pH of ∼0.1-0.3 units (total pH scale) by the end of the century, depending on the carbon dioxide emissions scenario (IPCC, 2019). In light of this, larvae released by the *P. acuta* colonies (n=9) were pooled and allocated, as described below, to one of three treatments of modified magnitude of diel fluctuations: Control conditions (present-day ambient temperature and pH), Mid conditions and High conditions (Table 1, Table S1) based on potential scenarios including the benthic organism feedbacks on biogeochemistry and residence time effects on temperatures in coastal embayments, specifically Kāne‘ohe Bay (C. P. Jury & Toonen, 2019).

**Table 1.** Carbonate chemistry summary table. Carbonate seawater chemistry parameters calculated from measured values of temperature, salinity, pH (total scale), and total alkalinity (TA). Values are shown as mean ± standard error. CO_2_ = carbonate, pCO_2_ = partial pressure of carbon dioxide, HCO_3_ = bicarbonate, CO_3_ = carbonate, DIC = dissolved inorganic carbon.

| Treatment | Salinity (psu) | Temperature (°C) | pH (Total) | CO<sub>2</sub> (μmol kg<sup>-1</sup>) | pCO<sub>2</sub> (μmol kg<sup>-1</sup>) | HCO<sub>3</sub> (μmol kg<sup>-1</sup>) | CO<sub>3</sub> (μmol kg<sup>-1</sup>) | DIC (μmol kg<sup>-1</sup>) | Total Alkalinity (μmol kg<sup>-1</sup>) | Aragonite Saturation State |
| --- | --- | --- | --- | --- | --- | --- | --- | --- | --- | --- |
| Control (n=12) | 35.02 ± 0.05 | 27.36 ± 0.06 | 7.89 ± 0.01 | 15.45 ± 0.23 | 578.66 ± 8.81 | 1778.71 ± 4.33 | 163.20 ± 1.68 | 1957.36 ± 3.06 | 2182.83 ± 1.61 | 2.62 ± 0.03 |
| Mid (n=12) | 34.97 ± 0.07 | 29.00 ± 0.06 | 7.71 ± 0.01 | 23.88 ± 0.43 | 929.85 ± 16.84 | 1881.94 ± 4.36 | 121.94 ± 1.65 | 2027.76 ± 3.3 | 2182.95 ± 1.68 | 1.98 ± 0.03 |
| High (n=12) | 35.04 ± 0.07 | 30.92 ± 0.06 | 7.51 ± 0.01 | 39.18 ± 0.69 | 1594.48 ± 28.45 | 1980.20 ± 3.33 | 85.35 ± 1.39 | 2104.73 ± 2.87 | 2190.15 ± 2.16 | 1.40 ± 0.02 |

Temperature and pH treatment conditions during the experiment were programmed to mimic the natural daily fluctuations at the collection site, which were monitored throughout the 20 days preceding the experiment (before introducing larvae) in the Control tank (Figure 1A, B). Experimental set-up consisted of flow-through 20-L polycarbonate tanks (n=3 per treatment, hereafter called larval tanks) within an outdoor mesocosm water bath (n=1 per treatment). Mesocosms were equipped with shade cloths as described above. Treatment conditions were generated in header tanks (120-L coolers) using a pH and temperature feedback system with a micro-processor controlled power strip (Apex Aquacontroller, Neptune Systems), and water delivered to each tank with a magnetic drive pump (Pondmaster Pond-mag Magnetic Drive Water Pump Model 5, Danner Manufacturing). To generate the pH treatments, two food-grade CO_2_ cylinders were connected to an automatic gas cylinder changeover system (Automatic Gas Changeover Eliminator Valves, Assurance Valve Systems). CO_2_ was added to the seawater in the header tank on-demand through gas flow solenoids (MA955, Milwaukee Instruments), based on the pH (NBS) reading of pH probes (Apex pH, Neptune Systems) in the larval tanks. CO_2_ was delivered via airlines with a venturi injector (Forfuture-go G1/2 Garden Irrigation Device Venturi Fertilizer Injector, BE-TOOL) connected to a water circulating pump (Pondmaster Pond-mag Magnetic Drive Water Pump Model 5, Danner Manufacturing). pH probes (Apex pH, Neptune Systems) were calibrated before and during the experiment using NSB pH calibration solutions (pH 7 and 10, NBS scale RYS377CN8B-1, Neptune Systems). During the experiment, pH was recorded every minute in each experimental tank through the Apex system. For temperature control, submersible heaters (GH-200W, AquaTop) were placed in the header tanks and the larval tanks (one heater per larval tank and header tank) to increase the temperature in the Mid and High temperature treatments, based on the temperature probes of the Apex System located in the larval tanks. During the experiment, temperature (°C) was recorded every 60 seconds in each experimental tank through the Apex system, as well as using underwater loggers (Hobo Pendant Water Temp/Light MX2202, Onset Computer Corp) placed at the same height as the settlement chambers (one HOBO logger per larval tank) at 5 minutes intervals. The same underwater loggers were also used to record light (lux) measurements every 5 minutes (Supplementary Figure 1B).

**Figure 1.**
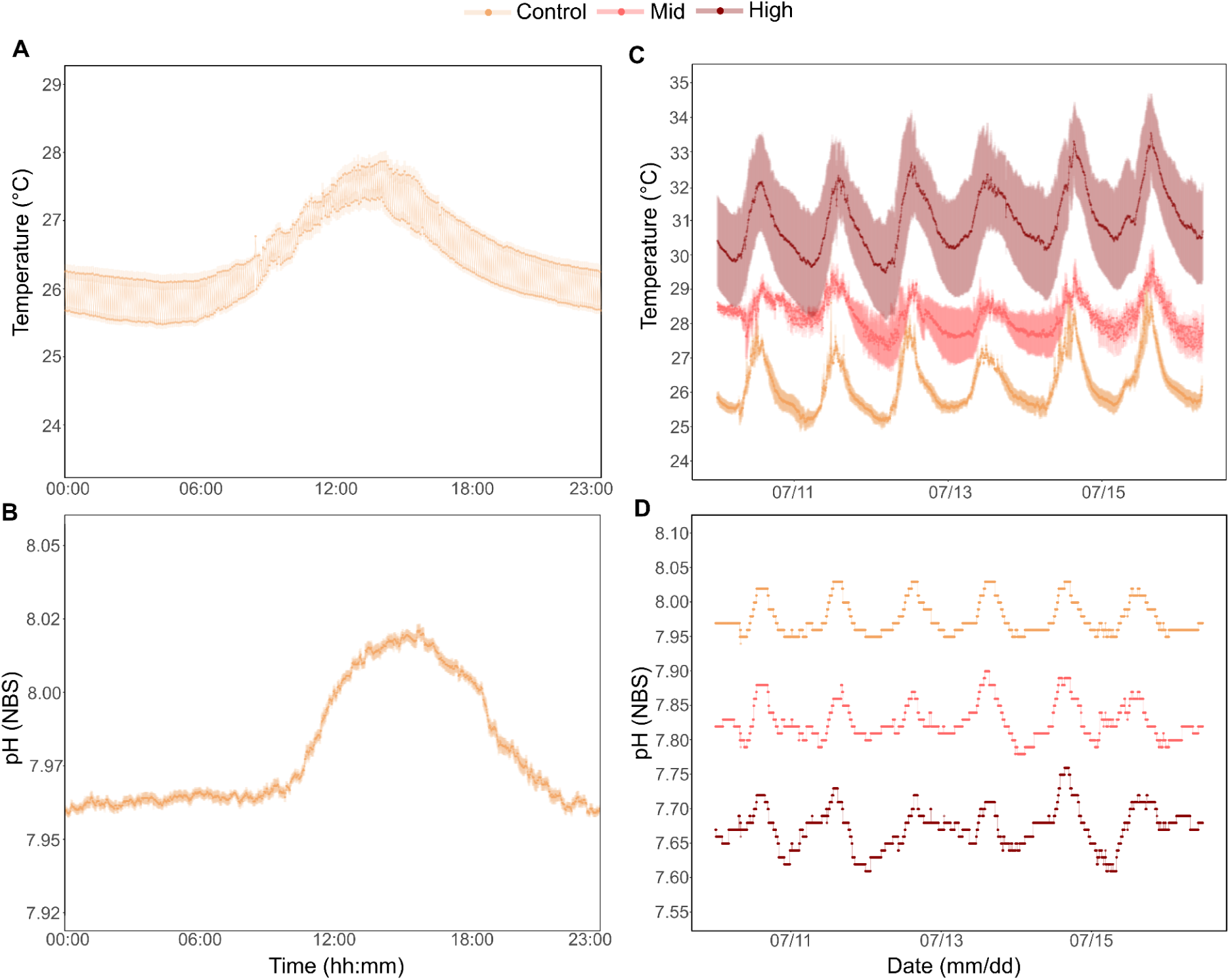
Environmental variability monitored via Apex Aquacontroller (Neptune Systems) before and during the experimental period. Mean daily temperature (A) and pH (B) monitored in the Control tank over the 20 days preceding the experiment (before introducing larvae). Mean temperature (C) and pH (D) values monitored during the experiment in the Control, Mid, and High larval tanks. Data is shown as means (solid lines and dots) ± sem (shaded lines).

Daily ambient temperature and pH fluctuated between ∼25.5-27.9°C, and ∼7.96-8.03 (pH NBS scale), respectively (measured as described above using HOBO loggers, accuracy ± 0.53°C, and the Apex pH probes from the Neptune system, accuracy ± 0.01 pH NBS scale; Figure 1A, B). Fluctuating diurnal temperature levels during the experiment were maintained between 26.9-29.5°C and 29.3-33.3°C in the Mid and High treatments, respectively (Figure 1C), and pH levels were maintained between 7.78-7.90 (pH NBS scale) and 7.61-7.76 (pH NBS scale) in the Mid and High treatments, respectively (Figure 1D).

### Seawater carbonate chemistry

Discrete measurements of pH (mV; Mettler Toledo 212 InLab Expert Pro pH probe and Orion Star A series A325, Thermo Fisher Scientific), salinity (conductivity psu; Orion DuraProbe 4-Electrode Conductivity Cell and Orion Star A series A325, Thermo Fisher Scientific), and temperature (°C, digital thermometer 4000EA, Traceable® Products) were taken up to 3 times a day throughout the duration of the experiment in each tank. Calibration curves of the relationship between mV and temperature in a tris standard (Dickson Laboratory Tris Batch T27 Bottles 269, 236 and Batch T26 Bottle 198) were conducted before and during the experiment. Water samples for carbonate chemistry were collected in 250 mL dark plastic bottles from each larval tank at four time intervals during the experiment: 10th, 12th, 14th, and 16th of July (n=3 samples per treatment per time interval; n=12 total samples per treatment). Samples were collected between 12 to 12:30 pm each day. Water samples were poisoned with 0.05% mercuric chloride within 30 minutes of collection to preserve the carbonate chemistry. Total alkalinity (TA) was determined using an open cell titration method (SOP3b; (Dickson et al., 2007)) with a certified HCl titrant (Dickson lab). TA measurements exhibited <1% error when compared against certified reference materials (Dickson lab CRM Batch 178). Daily measurements of temperature and salinity, along with TA values, were used to calculate total pH, as well as levels of carbon dioxide (CO_2_), partial pressure of carbon dioxide (pCO_2_), bicarbonate (HCO_3_ ^-^), carbonate (CO_3_ ^2-^), dissolved inorganic carbon (DIC), and aragonite saturation (Ω_Aragonite_) in the seawater using the seacarb package (v3.2; Gattuso et al. 2018) in R (v4.3.1) and RStudio (v2023.06.2; R Core Team 2023) with the following parameters: calculated total pH, calculated total alkalinity, flag=8 [pH and ALK given], P=0, Pt=0, Sit=0, pHscale=“T”, kf=“pf” (Perez & Fraga, 1987), k1k2=“l” (Lueker et al., 2000), ks=“d” (Dickson 1990).

### Primary polyps sampling

On the day of larval release (July 10th, 2022), pools of larvae (n=25 larvae per chamber) were allocated to square settlement plastic chambers (118 mL square shaped Ziploc Food Storage Meal Prep Containers) with an open top 153 µm mesh on the bottom and three sides, to allow for water exchange, and floated inside the larval tanks. Two aragonite plugs, with waterproof paper (1cm x 1cm; Rite In The Rain Waterproof DURARITE Copier Paper) attached on top to allow for easy and non-damaging removal of settled polyps, were placed in each chamber to provide a settlement substrate for the larvae. Prior to the addition of the larvae, the chambers were conditioned in ambient seawater in each corresponding tank during the 20 days preceding the experiment. After 6 days of treatment exposure on 16 July 2022, survival and settlement rates were assessed by counting the number of swimming larvae and settled primary polyps in the settlement chambers. Survival rates were calculated by dividing the number of swimming larvae and settled primary polyps over the total number of larvae that were allocated to the chambers (n=25 larvae chamber^-1^). Settlement rates were calculated by dividing the number of settled individuals over the total number of larvae originally allocated to the chambers, and settlement capacity of the surviving individuals were calculated by dividing the number of settlers over the number of larvae that remained alive per each treatment. Primary polyps were sampled by gently detaching them from the waterproof paper using a sterile scalpel, and were assigned to either molecular or skeletal analyses. For molecular analysis, primary polyps (n = 12 from each larval tank, n = 36 total per treatment) were preserved in 300 uL of DNA/RNA Shield (Zymo Cat# R1100-250, Zymo Research Corporation) and kept at -80°C until extraction. For skeletal analysis, primary polyps were put in 35 mm petri dishes (n = 3 per treatment, n = 3 polyps per dish) and stored with 80% ethanol until scanning electron microscopy imaging (SEM).

### Skeletal analysis

To determine if the main features characteristic of early skeleton formation (i.e., basal plate septa, and skeletal fibers arrangement) formed under all conditions, the tissue of the primary polyps was removed using 1% sodium hypochlorite (NaClO) for 10 minutes (n = 3 per treatment), rinsed using DI water, and then dried overnight. The skeletal samples were vacuum coated with 5 nm gold (for conductivity) and examined under a scanning electron microscope (SEM) ZEISS SigmaTM SEM (Germany), by using an in-lens detector (2 kV, WD = 3.9-4.9 mm), to assess presence or absence of the basal plates and septa, and the skeletal fibers arrangement.

### RNA extraction and sequencing

RNA was extracted from samples using the Zymo Quick-DNA/RNA Miniprep Plus (Zymo Cat# D7003, Zymo Research Corporation) kit following the manufacturer’s instructions. RNA quantity and quality were assessed with a Qubit 3.0 Fluorometer (Thermo Fisher Scientific) and an Agilent Tapestation 4200 (Agilent Technologies). RNA sequencing libraries were prepared using NEBNext Ultra II Directional RNA Library Prep Kit for Illumina (NEB Cat# E7760S, New England Biolabs) following the manufacturer’s instructions at Azenta Life Sciences. Libraries were sequenced at Azenta Life Sciences using an Illumina NovaSeq X in a 2x150 Paired End (PE) configuration.

### RNA-seq reads processing

Fastp (v0.19.7; (Chen et al., 2018)) was used to trim raw reads of adapters and poor quality sequences, and trimmed read quality was assessed using FastQC (v0.11.8, Java-1.8; Andrews 2010) and MultiQC (v1.9; (Ewels et al., 2016)). The *P. acuta* reference genome Version 2; (Stephens et al., 2022) predicted genes and proteins, and functional annotations were obtained from http://cyanophora.rutgers.edu/Pocillopora_acuta/. Trimmed reads were aligned to the reference genome using Hisat2 (v2.2.1; (Kim et al., 2019)). Aligned bam files were assembled to the reference genome using StringTie (v2.2.1; (Pertea et al., 2015)), and the StringTie prepDE python script was used to generate a gene count matrix (Pertea et al., 2015). Raw sequencing data is stored under NCBI BioProject PRJNA1107956.

### Differential expression and functional analysis

All expression and functional analyses were performed in R (v4.3.1) and RStudio (v2023.06.2; R Core Team 2023). Gene counts were filtered using pOverA from the package genefilter (v1.82.1), and genes were only retained for analysis if counts were greater than or equal to 5 in 25% of the samples (pOverA=0.25, 5; Gentleman et al. 2024). This threshold was chosen based on our experimental design of 12 total samples with a minimum of 3 samples per treatment group. The criteria ensured that genes were retained if they were expressed (count > 5) in at least 3 out of 12 samples (25%), allowing for potential differential expression between treatment groups, while removing genes with consistently low counts across all samples. Differential gene expression was assessed using DESeq2 (v1.40.2; design = ∼Treatment) with the Wald likelihood test ratio with a false discovery rate p-value adjustment (Love et al., 2014). Genes were considered differentially expressed if they had an adjusted p-value < 0.05. Differentially expressed genes were identified between pairwise treatment comparisons (i.e., High v. Control, High v. Mid, Mid v. Control).

GO enrichment analyses were performed with the R package GO-MWU (https://github.com/z0on/GO_MWU, which uses adaptive clustering of GO terms and Mann-Whitney U tests to determine overrepresented and underrepresented functions in the differentially expressed genes (FDR < 0.05; (Wright et al., 2015)). Log fold change was used as input for the Mann-Whitney U tests, which compares ranks among the log fold changes to assess whether the distribution of ranks of genes included in a GO term diverges significantly from that of other expressed genes. The deviation of a GO term from the rest of the data set is quantified by its delta rank (i.e., the difference in the mean rank of the GO term and the mean rank for all other GO terms; (Dixon et al., 2020). A positive delta rank indicates the GO term tends toward overrepresented, whereas a negative delta rank indicates the GO term tends toward underrepresented. The similarity of functional responses between treatment comparisons was compared by plotting the delta ranks of the treatment comparisons against one another. These plots do not represent a formal statistical test, as the data points (GO terms) often encompass overlapping sets of genes and thus are not independent (Dixon et al., 2020). However, the plots represent an effective way to compare functional similarity between the differential expression sets of the treatment comparisons.The tighter the positive relationship between delta ranks, the more similar the GO enrichment is for the differential expression sets of the treatment comparisons. To further support the relationship between the delta rank comparisons, a rank regression model was fit to the data to calculate the slope of the line using csranks (v1.2.3, https://github.com/danielwilhelm/R-CS-ranks; (Chetverikov & Wilhelm, 2024; Mogstad et al., 2024)). Biological Process (BP) and Molecular Function (MF) GO terms were analyzed using this approach.

### Biomineralization toolkit genes

A list of proteins and their associated sequences related to coral biomineralization was produced from *Stylophora pistillata*, a species in the same family (Pocilloporidae) as *P. acuta* (Scucchia, Malik, Putnam, et al., 2021; Scucchia, Malik, Zaslansky, et al., 2021). These protein sequences were compared to *P. acuta* protein sequences via BLASTp (v2.9.0; evalue 1E^-40^, max_target_seqs 1, max_hsps 1; (Altschul et al., 1990)). The putative *P. acuta* biomineralization hits were compared to the poverA filtered gene count matrix and the differentially expressed gene list to determine if any biomineralization toolkit-related genes were differentially expressed.

### Statistical analyses of environmental and performance data

Environmental parameters (temperature, salinity, total pH, CO_2_, pCO_2_, HCO_3_ ^-^, CO_3_ ^2-^, DIC, TA, and Ω_Aragonite_) were analyzed with a one-way analysis of variance (ANOVA) test with ‘Treatment’ as the independent variable. Treatment-specific differences were further assessed using Tukey post-hoc tests (p-value < 0.05). CO_2_ and pCO_2_ were log-transformed to fit ANOVA assumptions of normality. Salinity and TA were unable to be transformed to fit ANOVA normality assumptions; therefore, the non-parametric Kruskal-Wallis test with the Bonferroni correction was utilized, followed by the non-parametric Dunn test for post-hoc differences (p < 0.05).

Survival and settlement data were tested for normality using the Shapiro-Wilk test (normality assumed if p > 0.05) and confirmed with quantile-quantile plots, and for homogeneity of variance using the Levene test (p > 0.05). A one-way ANOVA was used to test differences between treatments, followed by Tukey test for post-hoc comparisons. Pairwise comparisons were considered significant if the post-hoc p < 0.05. Statistical tests were performed in R (v4.3.0) and RStudio (v2024.04.1).

## Results

### Carbonate chemistry

Temperature (°C), pH (total), CO_2_ (μmol kg^-1^), pCO_2_ (μmol kg^-1^), HCO_3_ (μmol kg^-1^), CO_3_ (μmol kg^-1^), DIC (μmol kg^-1^) and aragonite saturation (Ω_Aragonite_) all differed significantly by treatment (Figure S2; Table 1). While total alkalinity (μmol kg^-1^) in the High treatment was statistically higher than total alkalinity in the Control or Mid treatments (2190 v. 2182 µmol kg^-1^; Figure S2I; Table 1), the difference was lower than the 1% error from the CRM (∼20 µmol kg^-1^). Salinity was not significantly different across treatments. Further statistical details can be found in Table S1.

### Survival and settlement rates

Survivorship (Figure 2A) was significantly lower in corals from the High treatment compared to corals from the Control and the Mid treatments (ANOVA, F(2, 6) = 11.08, p < 0.01; Tukey’s multiple comparisons test, p = 0.009, p = 0.04, respectively). Similarly, percent settlement relative to total input of larvae (Figure 2B) were significantly lower in corals from the High treatment than rates in corals from the Control and the Mid treatments (ANOVA, F(2, 6) = 11.4, p < 0.01; Tukey’s multiple comparisons test, p = 0.01, p = 0.02, respectively). This was however due to mortality and therefore settlement capacity (percent settlement relative to the total surviving larvae; Figure 2C) was not significantly different across conditions (ANOVA, F(2, 6) = 3.68, p > 0.05; Tukey’s multiple comparisons test, p > 0.05).

**Figure 2.**
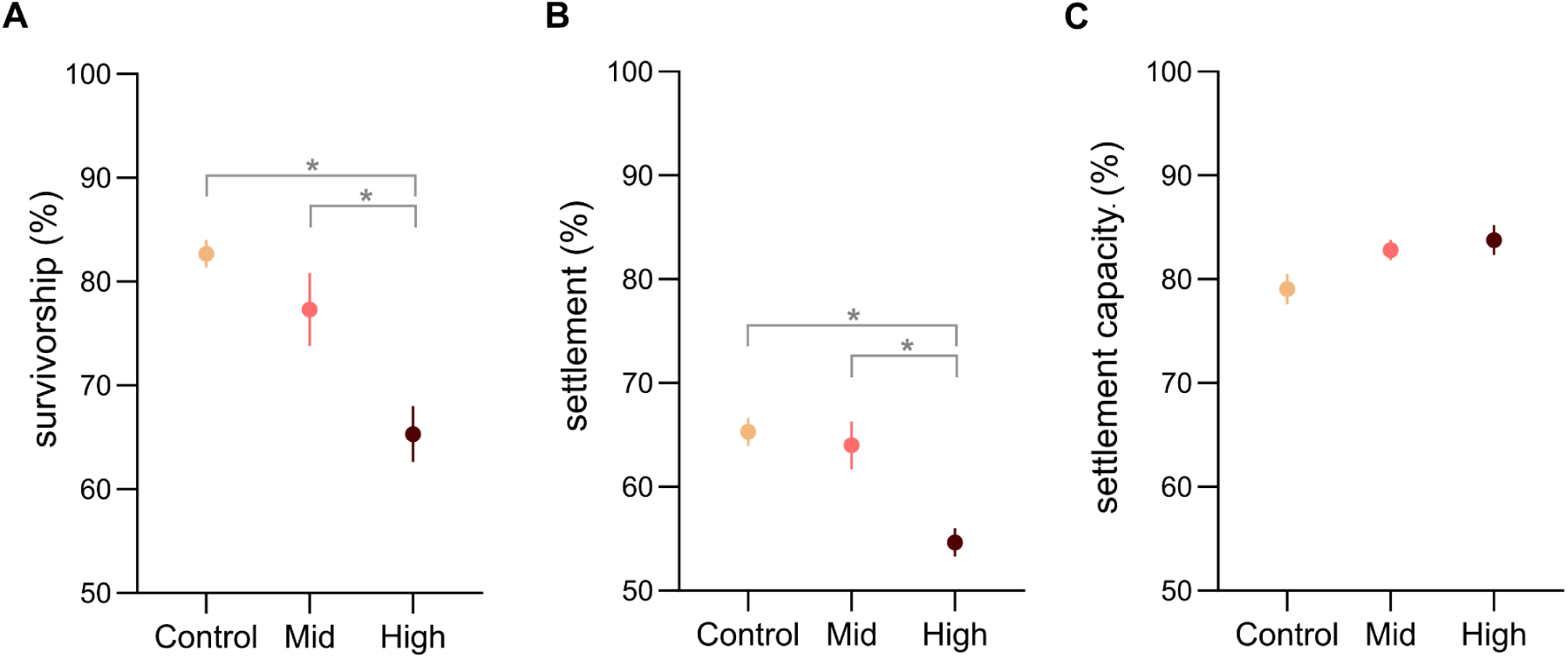
Survival and settlement of *P. acuta* across temperature and pH conditions. Mean number (as percentage; mean ± SEM) of (A) surviving individuals (swimming larvae + settled primary polyps relative to 25 larvae per chamber at the start of the experiment), (B) settled primary polyps (settled primary polyps relative to 25 larvae per chamber at the start of the experiment), and (C) settlement capacity (settled primary polyps relative to the swimming larvae + settled primary polyps alive) per treatment (Control, Mid, High) at the end of 6 days of experimental exposure. Asterisks (*) indicate statistical differences between conditions (n = 3 chambers per treatment; p-value < 0.05, One-Way ANOVA and Tukey’s multiple comparisons test).

### Skeletal features

SEM images of the primary polyps revealed the skeleton growth pattern under the different temperature and pH conditions. Deposition of the basal plate and primary and secondary septa was observed in all treatments (Figure 3A-C), and higher resolution images of the skeletal spines reveal similar granulated surfaces constituted of bundles of fibers (Figure 3D-F) in all treatments. At higher magnification, analogous packing of the rounded bundles is observable in all treatments, though only at the High treatment condition elongated needle-like aragonite fibers are visible in between the bundles (Figure 3G-I, yellow arrows in I).

**Figure 3.**
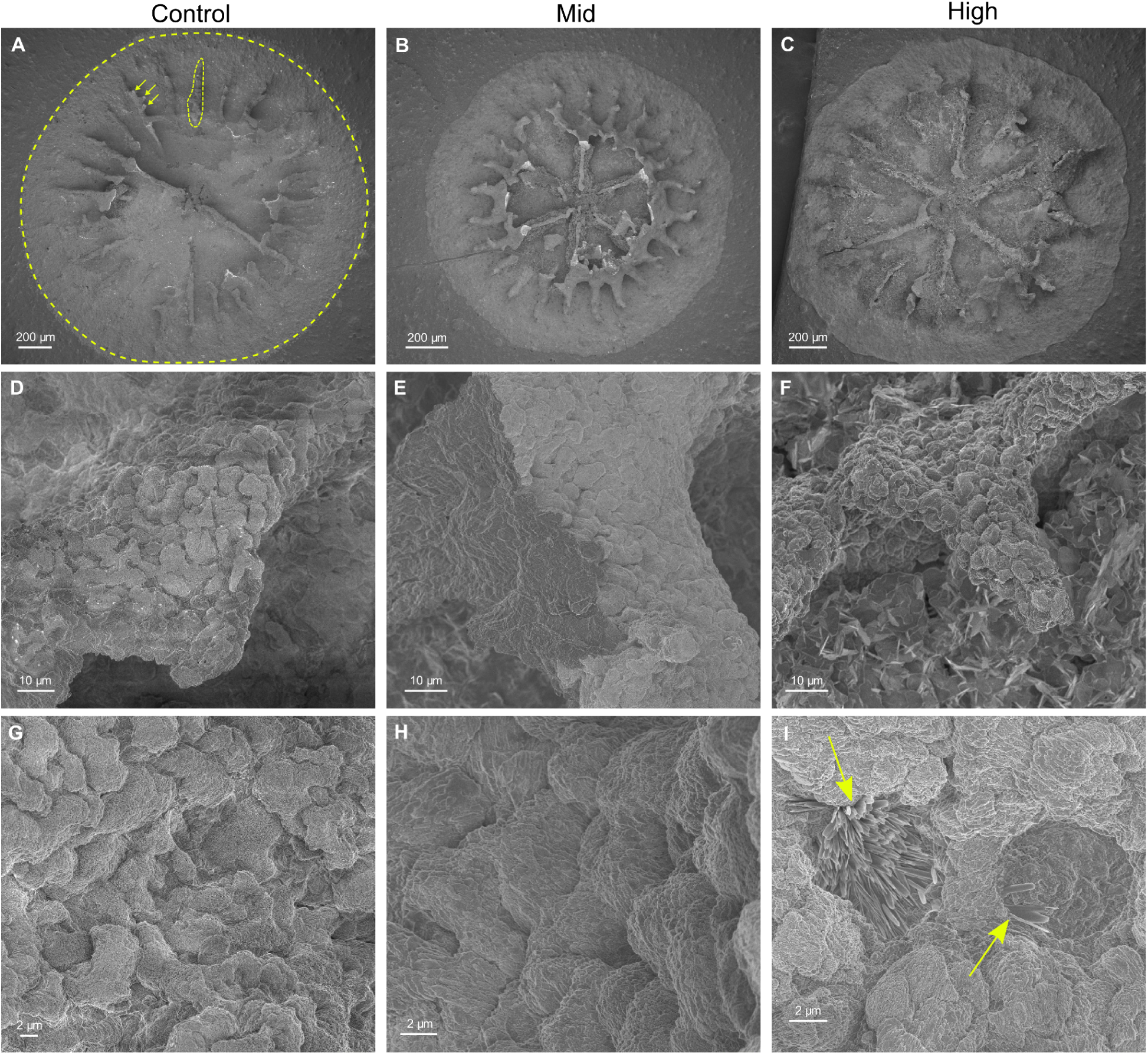
Structural features of *P. acuta* cultured at different temperature and pH conditions. Overall view of the primary polyps (A-C); example basal plate (yellow dotted-line circle), septum (yellow dotted line) and spines (yellow arrows) are shown in A. Enlargements of the spines on the skeleton septa (D-F), and enlargements of the spines surface showing details of the texture of the mineral bundles (G-I). The yellow arrows in I indicate the elongated needle-like fibers.

### Differential expression

RNA-Sequencing yielded an average of 29,892,892 total reads per sample (Table S2). After trimming, samples retained an average of 24,126,110 reads (Table S2) for a range of 79.8 to 81.5% of total reads retained after trimming (Table S2). Alignment with Hisat2 to the *P. acuta* genome yielded a range of 57.2 to 83.5% alignment and an average alignment of 78.6% across samples (Table S2).

24,185 genes out of 33,730 total genes remained for differential expression analysis after pOverA filtering of > 25% of the samples having greater than or equal to 5 reads. Comparisons between the High and Control treatments and the High and Mid treatments revealed 672 and 625 differentially expressed genes, respectively (Table S3). Notably, only 7 genes exhibited differential expression between the Mid and Control treatments (Table S3). These results suggest divergent molecular responses in primary polyps exposed to the High treatment compared with those in the Control and Mid treatments, as demonstrated in the High samples clustering together in the DEG heatmap (Figure 4A). Additionally, PCA of DEGs found that High treatment samples clustered away from the Mid and Control samples on the PC1 axis, accounting for 69% variance (Figure 4B).

**Figure 4.**
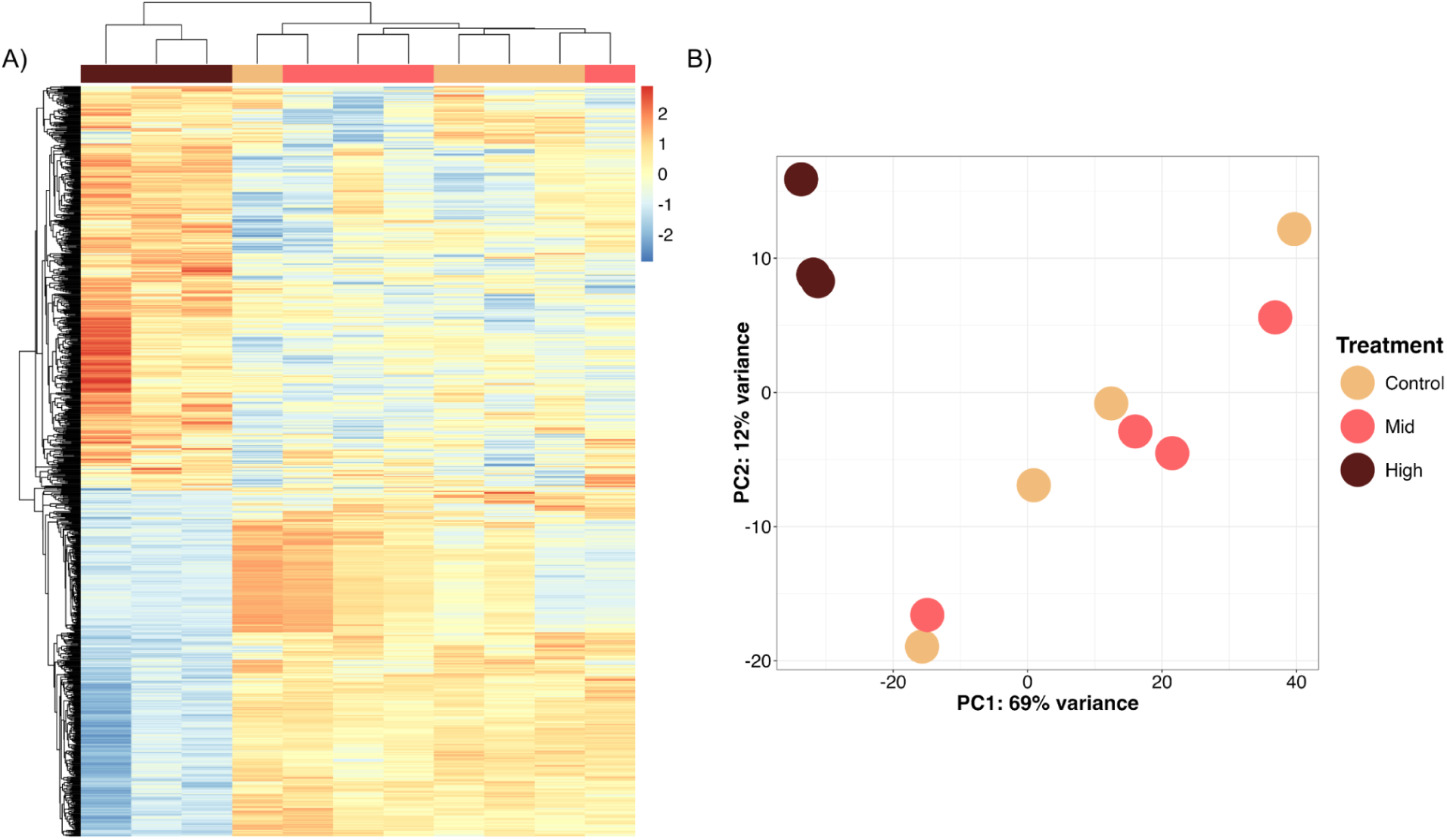
Differential gene expression patterns. Heatmap (A) and principal component analysis (PCA; B) of differentially expressed genes between treatments at the primary polyp stage. (A) Clusters of differentially expressed genes in the heatmap are grouped based on Euclidean distances calculated from their scaled expression profiles. Each row represents a gene, and the colors reflect the relative expression levels of the genes, scaled across samples to highlight differences between treatments. (B) The PCA visualizes sample-level variation in gene expression. It is based on sample-to-sample distances computed using the singular value decomposition technique, identifying principal components that explain major sources of variation between treatments.

There were only 4 downregulated genes and 3 upregulated genes in the Mid treatment compared to the Control treatment (Table S3). The High treatment had 306 downregulated and 366 upregulated genes relative to the Control treatment (Table S3). The High treatment had 307 downregulated and 318 upregulated genes relative to the Mid treatment (Table S3).

In total, 372 DEGs were shared between High v Control and High v Mid comparisons. 1 DEG was shared between Mid v Control and High v Control, 3 DEGs were shared between Mid v Control and High v Mid, and 1 DEG was shared between all comparisons. Among all comparisons, there were 926 unique differentially expressed genes across treatment comparisons (Table S4).

### Functional analysis

Given the large separation in gene expression between the cluster of Control and Mid samples and the High samples, we examined the consistency in functional differences in expression between High v. Control and High v. Mid by comparing the delta ranks (i.e., the difference in the mean rank of the GO term and the mean rank for all other GO terms; (G. Dixon et al., 2020))) of treatment comparisons. Terms in the positive quadrant of both the X and Y axes are significantly overrepresented in both comparisons, while terms in the negative quadrant of both the X and Y axes are significantly underrepresented in both comparisons. Terms falling closer to the 1:1 line indicate consistency in function in the comparisons being made.

Rank regression model estimated the slope of the BP comparison of delta ranks of High v. Control and High v. Mid to be 0.99 (Figure 5A; Table 2; Table S5) and no terms with delta ranks were present in the bottom right or top left quadrants of these comparisons. The top BP GO terms that showed high enrichment in the High treatment compared to both Mid and Control treatments included single strand break repair, G-quadruplex DNA unwinding, V(D)J recombination, negative regulation of telomere capping, and mRNA 3’-splice recognition (Figure 5A; Table 2; Table S5). The top underrepresented BP terms included the lipoxygenase pathway, axonemal dynein complex assembly, long-chain fatty acid biosynthetic process, fatty acid beta-oxidation, and other metabolic processes (tetrapyrrole, cellular aldehyde, NAD, cobalamin; Figure 5A; Table 2; Table S5).

**Figure 5.**
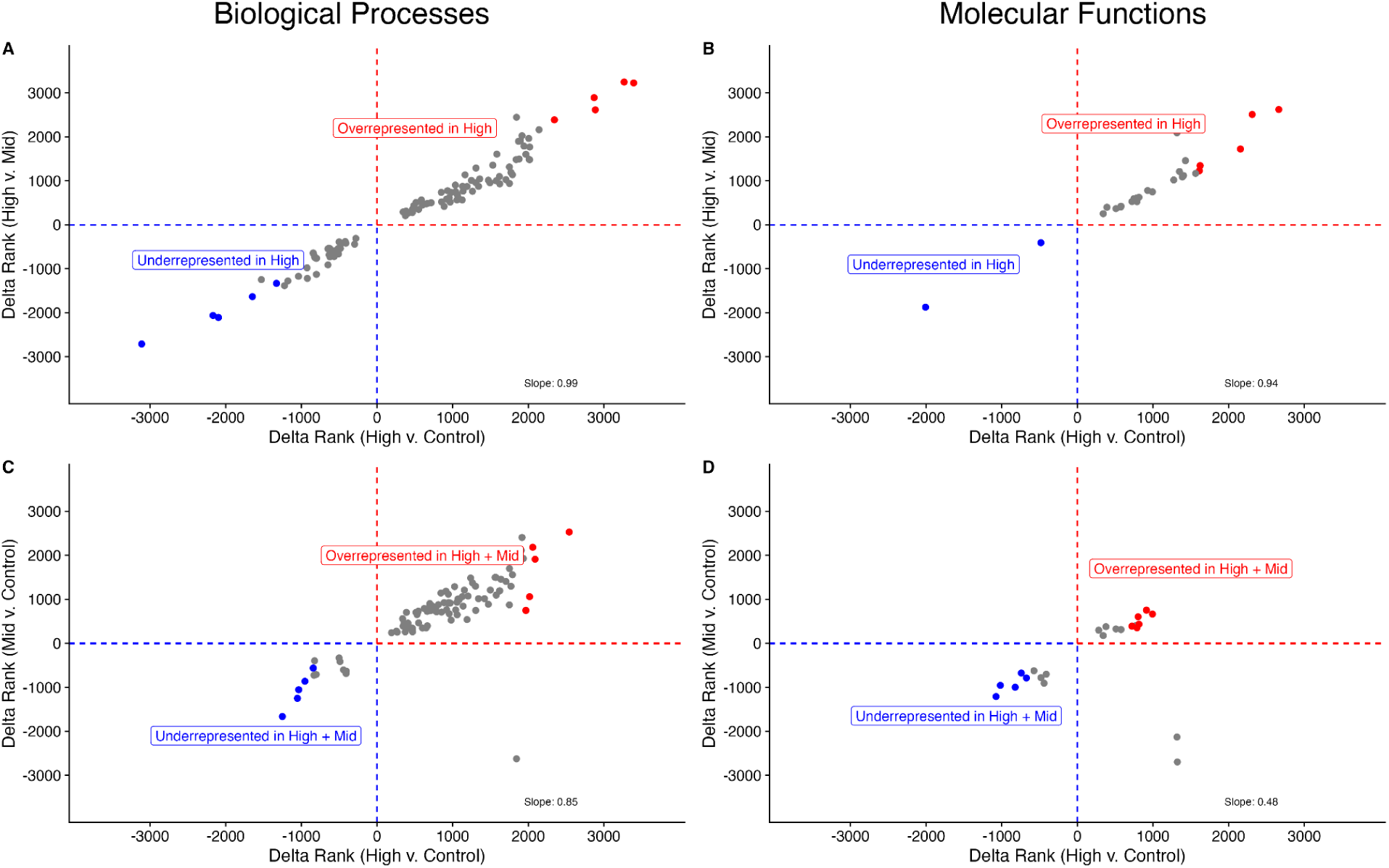
Delta ranks of (A) Biological Processes and (B) Molecular Function GO terms from the High v. Control (x-axis) and the Mid v. Control (y-axis) treatments; GO terms shown in color represent the top five overrepresented and underrepresented terms, as listed in Table 2. The delta ranks of (C) Biological Processes and (D) Molecular Function GO terms from the High v. Control (x-axis) and the High v. Mid (y-axis) treatments; GO terms shown in color represent the top five overrepresented and underrepresented terms, as listed in Table 3.

**Table 2.** Biological Processes and Molecular Function GO terms overrepresented and underrepresented in the High treatment relative to the Mid and Control treatments. Table corresponds to the colored points in Figure 5A, B.

|  | Ontology | GO | Term | Delta ranks (x, y) |
| --- | --- | --- | --- | --- |
| Overrepresented<br>in High relative to<br>Mid and Control | Biological Processes | GO:0044806 | G-quadruplex DNA unwinding | 3393, 3224 |
|  | Biological Processes | GO:0000012 | single strand break repair | 3267, 3245 |
|  | Biological Processes | GO:0033151 | V(D)J recombination | 2886, 2613 |
|  | Biological Processes | GO:1904354 | negative regulation of telomere capping | 2870, 2890 |
|  | Biological Processes | GO:0000389 | mRNA 3'-splice site recognition | 2345, 2385 |
|  | Molecular Functions | GO:0017116;GO:0033677 | single-stranded DNA helicase activity | 2663, 2620 |
|  | Molecular Functions | GO:0070259;GO:0017005 | tyrosyl-DNA phosphodiesterase activity | 2311, 2507 |
|  | Molecular Functions | GO:0035173 | histone kinase activity | 2156, 1722 |
|  | Molecular Functions | GO:0003697 | single-stranded DNA binding | 1623, 1344 |
|  | Molecular Functions | GO:0003678 | DNA helicase activity | 1613, 1228 |
| Underrepresented<br>in High relative to<br>Mid and Control | Biological Processes | GO:0019372 | lipoxygenase pathway | -3111, -2710 |
|  | Biological Processes | GO:0070286 | axonemal dynein complex assembly | -2168, -2062 |
|  | Biological Processes | GO:0042759 | long-chain fatty acid biosynthetic process | -2096, -2108 |
|  | Biological Processes | GO:0009235 | cobalamin metabolic process | -1647, -1634 |
|  | Biological Processes | GO:0019674 | NAD metabolic process | -1529, -1247 |
|  | Molecular Functions | GO:0016702;GO:0016701 | oxidoreductase activity, acting on single donors with incorporation of molecular oxygen | -2007, -1874 |
|  | Molecular Functions | GO:0016491 | oxidoreductase activity | -482, -408 |

**Table 3.**
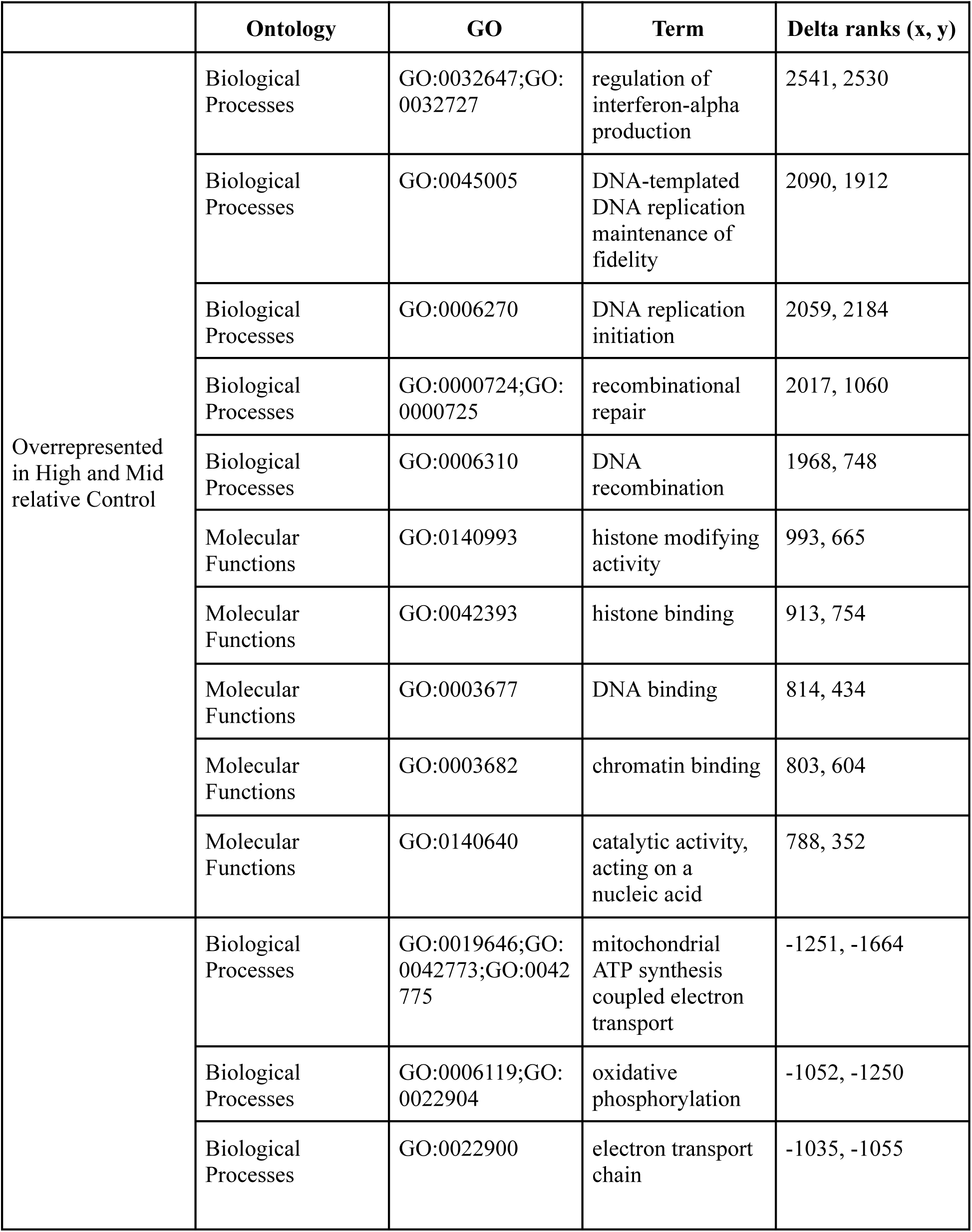
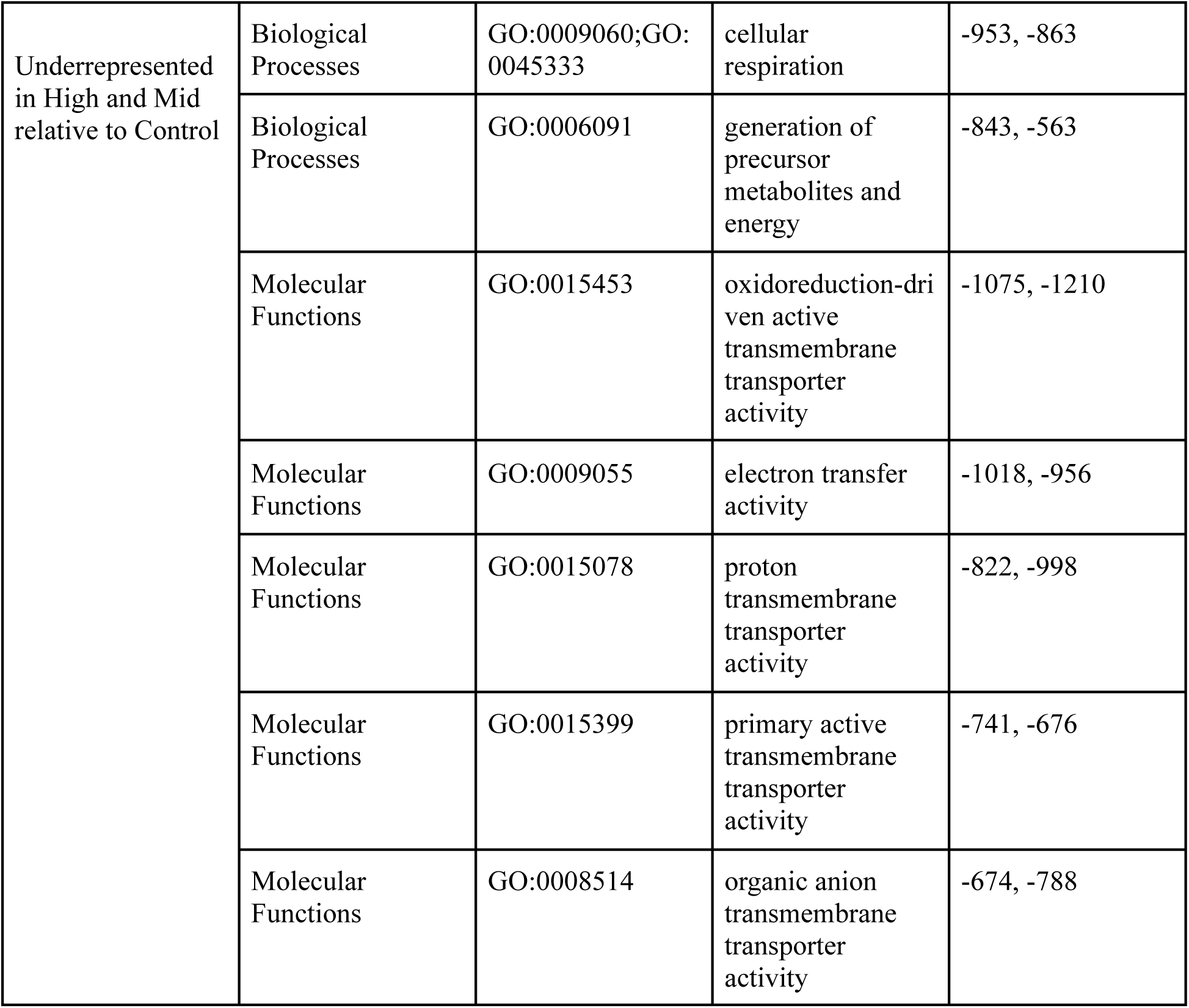
Biological Processes and Molecular Function GO terms overrepresented and underrepresented in the High and Mid treatments relative to the Control treatment. Table corresponds to the colored points in Figure 5C, D.

Similarly to BP, the slope of the MF comparison of delta ranks of High v. Control and High v. Mid was estimated to be 0.94. There were no terms with delta ranks in the bottom right or top left quadrants of this comparison. The top overrepresented MF terms in this context included single-stranded DNA helicase activity, DNA phosphodiesterase activity, histone kinase activity, single-stranded DNA binding, and DNA helicase activity (Figure 5B; Table 2; Table S5), whereas the top unrepresented MF terms included oxidoreductase activity and oxidoreductase activity, acting on single donors with incorporation of molecular oxygen (Figure 5B; Table 2; Table S5).

In contrast, when comparing the consistency in response of the High v. Control and the Mid v. Control for both BP and MP, the data points deviate further from the 1:1 line and some functions are even present in the bottom right quadrant, indicating high enrichment in one comparison, yet underrepresentation in the other comparison. The slope for the comparison of delta ranks in High v. Control and the Mid v. Control for BP was estimated as 0.85 by the rank regression model. The top overrepresented BP terms in the High and Mid treatment compared to the control included regulation of interferon-alpha production, DNA-templated DNA replication maintenance of fidelity, DNA replication initiation, recombinational repair, and DNA recombination (Figure 5C; Table 3; Table S5), and the top underrepresented BP terms included mitochondrial ATP synthesis coupled electron transport, oxidative phosphorylation, electron transport chain, cellular respiration, and generation of precursor metabolites and energy (Figure 5C; Table 3; Table S5).

The slope for the comparison of delta ranks in High v. Control and the Mid v. Control for was the lowest estimate at 0.48. The top overrepresented MF terms involved histone modifying activity, histone binding, DNA binding, chromatin binding, and catalytic activity, acting on a nucleic acid (Figure 5D; Table 3; Table S5). The top underrepresented MF terms included oxidoreduction-driven active transmembrane transporter activity, electron transfer activity, proton transmembrane transporter, primary active transmembrane transporter activity, and organic anion transmembrane transporter activity (Figure 5D; Table 3; Table S5).

### Biomineralization toolkit

Out of 172 protein sequences from the biomineralization toolkit, 168 of those sequences were in the filtered gene counts list, corresponding to 84 unique *P. acuta* genes (Table S6). Six biomineralization-related protein sequences, corresponding to five *P. acuta* DEGs, were identified as differentially expressed between treatments. Specifically, three and five DEGs corresponding to biomineralization processes were identified in the High v. Control and the High v. Mid comparisons, respectively (Table S7). There were no biomineralization-related DEGs identified in the Mid v. Control treatment comparison. There were two biomineralization genes that were differentially expressed in both the High v. Control and the High v. Mid comparisons. The first gene, Pocillopora_acuta_HIv2 TS.g23498.t1, which was downregulated in the High treatment relative to the Control and Mid treatments, was associated with functions related to alpha-collagen and co-adhesion (Table S7). The second gene, Pocillopora_acuta_HIv2 RNAseq.g7668.t1, was upregulated in the High v. Control and High v. Mid comparisons, but was uncharacterized (Table S7).

## Discussion

Temperature and pH vary daily within many reef habitats (Guadayol et al., 2014; Camp et al., 2018; Cyronak et al., 2020; De Almeida et al., 2023) and there is a strong relationship between an organism’s physiological limits and environmental fluctuations (Boyd et al., 2016; Rivest et al., 2017). Thus, experiments incorporating environmental variability into generated conditions are essential to fully capture the effects of multiple stressors on corals to global change (Boyd et al., 2016; Rivest et al., 2017). Our examinations unravel temperature and OA tolerance thresholds of *P. acuta* early life stages, which are able to withstand moderate warming and acidification conditions. However, more extreme conditions cause major reduction in survival, as well as notable changes in gene expression and skeletal growth.

### Environmental conditions outside tolerance thresholds decrease survivorship of P. acuta

Larval survivorship and successful settlement are critical bottlenecks for many marine organisms (Caley et al., 1996). Here, we found that larvae of *P. acuta* have the capacity for successful settlement under all treatment conditions, but the combination of higher temperature and pCO_2_ conditions under the High treatment led to significantly reduced survivorship and thus settlement rate (Figure 3). Similar to our results at moderate temperature increases and pH decreases (Mid), studies that have evaluated the effects of moderate temperature increases and pH decreases on brooding larvae have found that treatment did not affect coral survivorship and settlement, even in the case of longer incubation under treatment conditions compared to our study. For example, *Porites panamensis* planulae survival and settlement rates were not affected after 10 days under a combination of moderately higher temperature (+1-1.2°C) and lower pH (0.2-0.25 pH unit decrease) conditions compared to the control (Anlauf et al., 2011). Another study conducted on *P. acuta* recruits in Kāne‘ohe Bay did not find any significant difference in recruit abundance under moderately increased temperature (+2°C) and lower pH (0.2 pH unit decrease) compared to control conditions, over the course of a 22 month experiment (Bahr et al., 2020). Notably, *Seriatopora caliendrum* recruits had enhanced growth and survival when exposed to a diurnally oscillating pCO_2_ regime rather than a stable pCO_2_ regime, highlighting the importance of incorporating environmentally-relevant fluctuations into experiments (Dufault et al., 2012). However, survivorship and settlement of the spawning coral *Orbicella faveolata* larvae under a medium emission scenario (+1.5°C combined to -0.2 pH units) was reduced by ∼50% compared to the ambient treatment 7 days after spawning (Pitts et al., 2020).

In contrast to our observations of mortality and perturbed morphology and differential gene expression in *P. actua* spat in the High condition, extreme temperature and pCO₂ conditions (+2.4°C combined to -0.27 pH units) did not significantly affect survival rates of recruits of *Acropora intermedia* in a 33-day experiment (Sun et al., 2020), although exposure to warming and acidification conditions was carried out only after settlement in this study. Settlement and post-settlement survivorship of recruits of another *Acropora* species, *A. spicifera*, were also unaffected by extreme warming and acidification (+3°C combined to -0.47 pH units) in a 5-week experiment (Foster et al., 2015). Such diverse responses of corals to projected warming and acidification conditions may stem from experimental differences (e.g., exposure magnitude and variability, duration of the experiment), but also from differences in life-history traits (Boyd et al., 2016; Rivest et al., 2017; Torda et al., 2017), including reproductive strategy (i.e., broadcast spawner v. brooder) and mode of symbiont acquisition (i.e., horizontal v. vertical transmission). These life-history traits have important implications on how corals respond to environmental stress. For example, within the same reef location, larvae from horizontal broadcaster spawners (*Lobactis scutaria*) and vertical broadcast spawners (*Montipora capitata*) were both more susceptible to temperature and elevated nutrient content compared to larvae from a brooding vertical species (*P. acuta*; (Kitchen et al., 2020).

The relative resistance of brooded larvae, regardless of their symbiont transmission mode, may be due to parental investment. Larvae produced through broadcasting spawning typically have lower lipid energy reserves than larvae produced by brooding (40-60% and 85% dry weight, respectively; (Harii et al., 2007)), providing brooded larvae with more resources to dedicate to growth and metamorphosis while combating environmental stress. Vertically-transmitted symbionts may help the offspring survive and metamorphose, as symbiont-derived nutritional products ramp-up quickly post-fertilization, supporting larval growth and subsequent metamorphosis (Huffmyer et al., 2023). However, energetic demand rises during metamorphosis (Harii et al., 2002, 2010; Edmunds et al., 2013), coinciding with a reduction in the photosynthesis:respiration ratio during this delicate phase (Huffmyer et al., 2023). As such, decline in symbiont-derived nutrition driven by environmental stress could further exacerbate the energetic vulnerability of coral recruitment (Nakamura et al., 2011; Edmunds, 2023). In the High treatment, energetic resources for *P. acuta* recruits may not have been sufficient to sustain development, which becomes more energetically demanding under climate change (Edmunds et al., 2005, 2013), leading to significantly reduced survivorship (Figure 3). These differences, in conjunction with experimental methodologies, provide critical context to understanding how different coral species may respond to climate change.

### Biomineralization machinery function is maintained, although skeletal deposition may occur at slower rate under extreme conditions

The septa and basal plate, the two dominant and well characterized features of coral primary polyps (Neder et al., 2019; Mor Khalifa et al., 2021; Sugiura et al., 2021), formed in all treatments (Figure 3 A-C) without apparent deformities. Only in the High treatment, we observed needle-like fibers that appeared to have recently started to fill the pores in between the granular particle bundles (Figure 3 G-I). Initial skeleton building at the primary polyp stage includes the formation of the septa growing upward on the basal plate (Neder et al., 2019; Sugiura et al., 2021). Each septum consists of a central region called center of calcification (CoC) or rapid accretion deposits (RADs) characterized by a granular texture, and by elongated needle-shaped crystals arranged around the RADs (Cohen & McConnaughey, 2003; Von Euw et al., 2017). These needle-shaped aragonite crystals gradually fill the space to form the septum structure, and accumulate becoming the most abundant mineral material in the coral (Neder et al., 2019). The presence of these needle-like fibers only in the corals from the High condition and not in the other treatments (Figure 3 G-I) suggests that these corals were growing at a slower rate than the Control and Mid corals, with more time needed for the needle-like fibers to fill the spaces in the growing septa. On the contrary, the presence of more fibers could be a sign of more rapid growth at the High condition, as previous work has shown enhanced calcification under fluctuating pH conditions in coral recruits (Dufault et al., 2012). However, our gene expression data indicates that corals at the High conditions have depressed metabolism and reduced ATP production (Figure 5A). Considering that the calcification process becomes more energetically demanding under OA (Ries, 2011), our combined morphological and gene expression results suggest slower rather than faster calcification at the High condition compared to the other treatments.

Other studies have found that low pH alone slows down the skeleton growth and causes skeletal deformities in coral recruits (Foster et al., 2015; Scucchia, Malik, Zaslansky, et al., 2021). However, higher temperature seems to mitigate the negative effects of lower pH in both young corals and adults (Foster et al., 2015; Allison et al., 2022; Cameron et al., 2022). Such studies have been conducted with longer incubations under experimental conditions, suggesting that a shorter incubation time, as in our experiment, may not have revealed skeletal changes that potentially occur at later stages. Additionally, they suggest that the positive effect that higher temperature has on the calcification rate may be offsetting the negative effect of OA. Nevertheless, such beneficial effect is likely lost at temperatures that go beyond coral thermal optima, causing declines to calcification (Foster et al., 2014; Scheufen et al., 2017).

While in general slower coral skeletal growth has been attributed to increased energetic demands of the calcification process under OA, as corals require more energy to remove from the calcifying space the excess H⁺ produced during calcification (Ries, 2011), and specific disruption of the disruption of the function or activity of calcification-related genes is a growing area of study (Yuan et al., 2018; Bernardet et al., 2019; Scucchia, Malik, Zaslansky, et al., 2021; Yuyama et al., 2022). Looking at a set of biomineralization-related genes (Table S6) expressed here in *P. acuta* primary polyps (Table S7), we only found five differentially expressed genes in the High treatment relative to both the Control and Mid treatments. Among these, the alpha-collagen gene was downregulated in corals from the High treatment compared to corals from the Control and Mid treatments (Table S6). Collagen is an adhesion protein responsible for the attachment and binding of aragonite crystals along with other calcium-binding and adhesion proteins (Drake et al., 2013; Mummadisetti et al., 2021). The expression of biomineralization-related collagen gene has been found to show no differential expression in the gorgonian octocoral *Pinnigorgia flava* under the exposure to thermal and OA stress (Vargas et al., 2022) but to be suppressed in adults of the stony coral *Galaxea fascicularis* under OA (Lin et al., 2022). In the latter study, OA was found to induce concurrent changes in calcification and photosynthetic processes by altering the reallocation of dissolved inorganic carbon between the coral host and the symbionts, and to alter nutrient metabolic cycling which fuels calcification in the coral-algal symbiotic system. To date, no studies have evaluated how the synergistic effects of warming and OA specifically affect the expression of biomineralization genes in newly settled coral recruits and juveniles. Therefore, there is a critical need for studies identifying transcriptional changes underlying biomineralization mechanisms capturing multiple stages of coral development under multi-stressor scenarios. This would identify if skeletal growth is impacted at later developmental stages, and which gene expression changes may underlie such disruption in the coral host and/or the algal symbiont.

### Transcriptomic evidence for metabolic disruption prevalent in recruits exposed to High conditions

We found that in comparison to Control and Mid corals, *P. acuta* recruits from the High treatment exhibited downregulation of genes encoding for various ATP-consuming metabolic processes, such as tetrapyrrole, cellular aldehyde, NAD, carbohydrate derivative, lipids, and fatty acid metabolism (Figure 5A). Carbohydrates, lipids, and fatty acid are the main components of photosynthates (i.e., photosynthetically-derived products translocated from symbiont to coral host) (Gordon & Leggat, 2010; Burriesci et al., 2012), and the downregulation of related metabolic processes in High treatment recruits strongly suggests a reduction in symbiont-derived nutrition. This has been previously observed in corals exposed to elevated temperatures, where thermal stress disrupts symbiont functionality and nutrient transfer (Sun et al., 2020; Tremblay et al., 2016; Rädecker et al., 2021; Allen-Waller & Barott, 2023). Alternatively, or perhaps concurrently, environmental changes exceeding the stress-tolerance window of coral recruits, such as those conditions in the High treatment, may affect the optimal allocation of energy by reducing organismal capacities for assimilation and conversion of energy. Stress-induced increases in physiological maintenance costs divert resources to protective and reparative mechanisms, reducing capacities for energy assimilation and metabolic efficiency (Sokolova, 2013). In line with this, we observed upregulation of DNA damage-repair processes in the High treatment recruits (Figure 5A, B), a response which aligns with previous reports of corals activating similar pathways under thermal or OA stress (DeSalvo et al., 2010; Kaniewska et al., 2015; Dilworth et al., 2024).

Depressed metabolic states under high temperature and/or low pH conditions have been documented, both in adult corals (Kaniewska et al., 2012; Innis et al., 2021; Scucchia, Malik, Putnam, et al., 2021)) and during early life stages (Albright & Langdon, 2011; Nakamura et al., 2011; Cumbo et al., 2013; Putnam et al., 2013; Rivest & Hofmann, 2014; Moya et al., 2015; Scucchia, Malik, Zaslansky, et al., 2021). Metabolic suppression is considered an adaptive strategy to extend the time that an organism can survive during periods of environmental stress (Guppy, 2004; Sokolova, 2013). This hypometabolic state reduces capacities for assimilation and conversion of energy by suppressing the production and consumption of ATP, and non-essential functions such as growth and reproduction are halted (Sokolova & Pörtner, 2001; Marshall & McQuaid, 2011). Although metabolic depression can be important for promoting survival under stress in the short term, it can have long-term negative consequences for coral energetic status (Grottoli et al., 2006; Anthony et al., 2009) and ecological success (Anthony et al., 2008). For example, decreased coral metabolism and reduced energy reserves during heat stress or under OA conditions can lead to sublethal reductions in feeding (Ferrier-Pagès et al., 2010), reproduction (Levitan et al., 2014; Johnston et al., 2020; Leinbach et al., 2021; Bouwmeester et al., 2023), and skeleton growth (Albright & Langdon, 2011; Scucchia, Malik, Zaslansky, et al., 2021), which are all critical processes for species survival. Thus, under extreme conditions, *P. acuta* long-term species recruitment, survival, and overall fitness may be impaired.

We also observed downregulation of oxidoreductase-encoding genes in High treatment corals relative to corals in Control or Mid treatments, providing further evidence for compromised metabolic pathways in the High treatment corals (Figure 5B). Oxidoreductases are key enzymes in metabolic pathways such as glycolysis and oxidative phosphorylation, which drive ATP production and cellular energy generation (Das & Sen, 2024). Previous studies have found upregulation of oxidoreductases in corals under thermal stress (Rosic et al., 2014; Dixon et al., 2015), suggesting a compensatory mechanism to meet increased energy demands. In contrast, the downregulation of oxidoreductases observed here indicates that the synergistic and fluctuating stresses imposed in the High treatment elicited a different response, suppressing rather than enhancing these critical metabolic pathways. The suppression of oxidoreductases further underscores the compromised metabolic state of coral recruits under multiple stressors, potentially exacerbating energy deficits and reducing resilience to environmental perturbations.

### Environmental history shapes organismal stress-tolerance window, priming young coral stages to withstand Mid treatment conditions

In comparison to Control corals, corals from the Mid and High treatments downregulated cellular energy-regulating processes, including mitochondrial ATP synthesis coupled electron transport, oxidative phosphorylation, electron transport chain, cellular respiration, and generation of precursor metabolites and energy, as well as downregulation of genes encoding for transport activity, including those involved in ion transport and homeostasis. This downregulation implies alterations in ion gradients and transport mechanisms, which are indeed negatively affected by acidification in marine organisms leading to disruption of cellular homeostasis (Stillman & Paganini, 2015). Ion gradients across membranes are fundamental to driving numerous biological processes, including ATP production, the primary energy currency of the cell (Bonora et al., 2012). The observation of concurrent downregulation of ATP production processes suggests onset of a metabolic depression state; however, only High treatment corals exhibited significantly lower survival (Figure 2), as well as downregulation of ATP-consuming metabolic processes (tetrapyrrole, cellular aldehyde, NAD, carbohydrate derivative, lipids, and fatty acid metabolism). This disparity suggests that in Mid treatment corals, the pool of available metabolic energy is reduced but remains positive, thereby mediating survival. As the environmental variables further deviate from the stress-tolerance window, as in the High treatment, the available metabolic energy of the organism continues to decline until it reaches the threshold, at the expense of growth, reproduction, and activity (Sokolova, 2013). It is also possible that the full extent of exposure to the Mid treatment only becomes apparent with prolonged exposure, even though a previous study found that size and abundance of *P. acuta* juveniles in Kāneʻohe Bay was not impacted by combined ocean warming and acidification over a 22 month experiment (Bahr et al., 2020). However, health and survival of the recruits in the Bahr et al. (2020) study were negatively affected during the warmest month (October, our study was conducted in July), further indicating that environmental variations exceeding the width of the organism stress-tolerance window lead to negative consequences for coral fitness.

The width of the stress-tolerance window can be shifted by environmental history, the set of abiotic conditions to which individuals are acclimatized or adapted (Boyd et al., 2016; Rivest et al., 2017; Torda et al., 2017). Environmental history can influence stress tolerance by, for instance, transgenerational acclimatization (Putnam, 2021), which occurs when parents integrate information from their environment and affect the phenotype of their offspring (Byrne et al., 2020). Such parental effects can facilitate phenotypic plasticity, modulating individual responses and acting as a buffer towards environmental change (Gibbin et al., 2015; Putnam & Gates, 2015; Bonamour et al., 2019). Local adaptation of populations is another mechanism by which environmental history can shape stress tolerance (Rivest et al., 2017; Torda et al., 2017; Drury, 2020), acting by changes in the genome in response to selective pressures over multiple generations (Sanford & Kelly, 2011). Locations where daily maximum temperature and pH minima already reach levels predicted to occur in future climate scenarios might harbor populations possessing higher tolerance thresholds for environmental stress (Oliver & Palumbi, 2011; Schoepf et al., 2015; Camp et al., 2016; Rivest et al., 2017; Kirk et al., 2018; Thomas et al., 2018; Tanvet et al., 2023; Brown et al., 2024). Likewise, wide environmental fluctuations experienced by *P. acuta* corals in Kāne‘ohe Bay (Figure 1A, B) may prime offspring to endure moderate changes in temperature and pH (Putnam & Gates, 2015; Putnam et al., 2020). However, when climate-change-driven environmental changes break outside the natural variability window, such as under the High treatment, the organismal physiological threshold is reached, negatively affecting larval survival and settlement, with negative implications for cellular and organismal metabolism.

## Conclusion

The present study provides insights into the response of the early life stages of an abundant Pacific coral species to future projected warming and acidification scenarios, emphasizing the importance of considering environmental fluctuations to better understand how natural variability shapes organismal physiological thresholds and stress responses. While a higher number of replicates and sampling time points (i.e. older recruits) would clarify the physiological long-term costs of developing under future climate scenarios, this work provides evidence that corals inhabiting locations with large environmental fluctuations may be able to successfully recruit and grow under moderate environmental change. This potential for transgenerational “environmental memory” can elicit coral stress hardening (Putnam et al., 2017, 2020; Hackerott et al., 2021; Brown et al., 2022; DeMerlis et al., 2022) which may be utilized as proposed restoration technique (Van Oppen et al., 2015). However, it is clear that there is species-specific susceptibility to stress, even among species inhabiting the same location, due to factors including differences in symbiont and host physiological properties (Gibbin et al., 2015; Putnam et al., 2016; Strand et al., 2024). Furthermore, as warming and OA intensify, coral physiology will be pushed closer to its limits, and the energy required to sustain tolerance may narrow the intensity and duration of environmental stress that an organism can endure (Frieder et al., 2014). For stress-hardened corals, in fact, biological trade-offs for adaptation to extreme environments may come with compromises in genetic and energetic mechanisms and skeletal properties (Scucchia et al., 2023). Therefore, further determination of the extent and heritability of physiological tolerance across developmental stages and through multi-year studies is warranted to disentangle how environmental history will shape coral fitness across generations under future climate scenarios.

## Acknowledgments

We thank the Hawai‘i Institute of Marine Biology facilities staff and the Coral Resilience Lab for hosting us on Moku o Lo‘e and for their assistance in conducting the experiment. As guests, we recognize and give thanks for the land and water resources of the ʻāina and the traditional owners of the land, kānaka ʻōiwi, both past and present, as well as future generations, on which this experimental work was conducted in the Kāneʻohe Ahupuaʻa and the islands of Hawaiʻi. This work was supported by the Israel Academy of Sciences and Humanities (Aharon and Ephraim Katzir Study Grant to FS), the National Science Foundation (Graduate Research Fellowship to JA), and the US-Israel Binational Science Foundation (Award 2016321 to TM and HMP).

## Data Accessibility

Raw sequencing data is stored under the NCBI BioProject PRJNA1107956. All scripts and files employed to analyze the data are accessible through the electronic notebook https://github.com/fscucchia/Pacu_Warm_OA_Hawaii.

## Author Contributions

Funding: HP, TM; Experimental Design: FS; Field work: FS, JA, AH; Molecular work: FS, JA; Analyses: JA, FS; Writing: JA, FS; Editing: JA, FS, AH, HP, TM.

