## Supplemental Material for "Exposure to a gradient of warming and acidification highlights physiological, molecular, and skeletal tolerance thresholds in *Pocillopora acuta* recruits"

* equal contribution to authorship
### corresponding author

#### Supplemental Materal

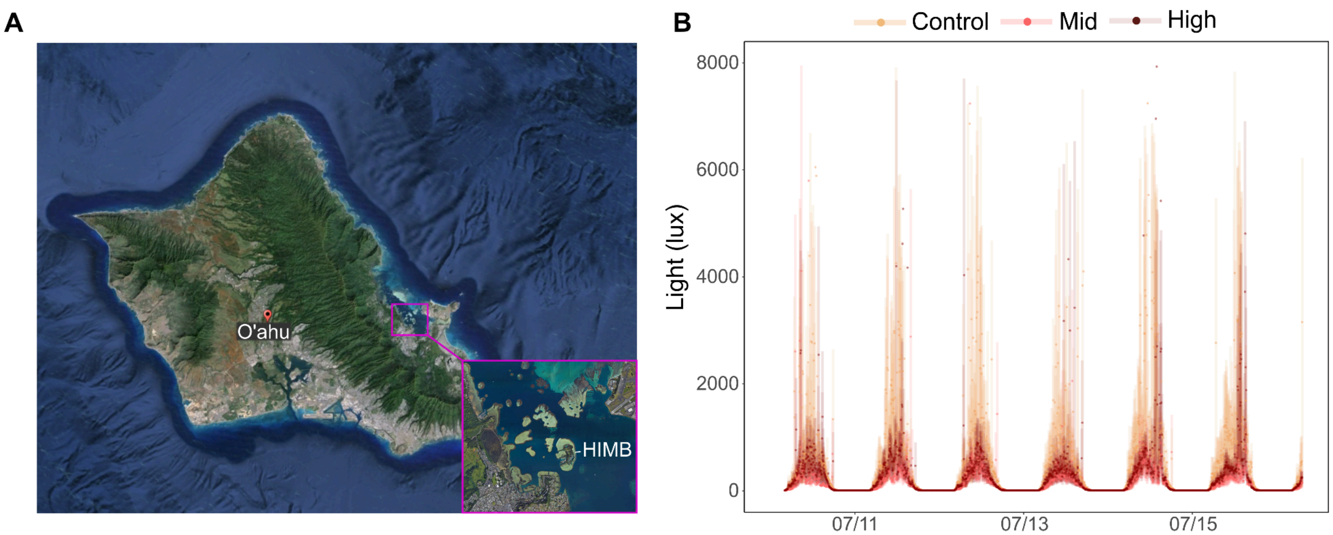


Figure S1. **Study site and light intensity monitored during the experimental period.** (A) Overview of O‘ahu, Hawai‘i, with a highlight box showing the study site. Mean daily light intensity (B) monitored during the experiment in the Control, Mid and High tanks. Data is shown as means (dots; n = 3 HOBOs per treatment) ± standard deviation (shaded lines).

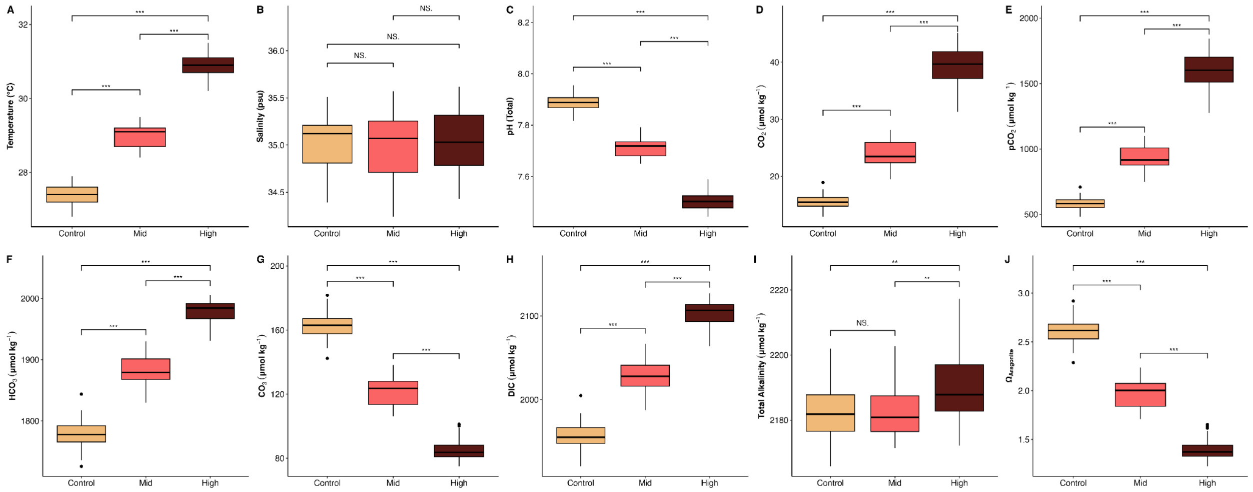


Figure S2: Environmental parameters from each treatment (Control, Mid, High). (A) Temperature (°C), (B) Salinity (psu), (C) Total pH, (D) $CO_2$ (μmol $kg^{-1}$), (E) $pCO_2$ (μmol $kg^{-1}$), (F) $HCO_3^-$ (μmol $kg^{-1}$), (G) $CO_3^{-2}$ (μmol $kg^{-1}$), (H) dissolved inorganic carbon (DIC; μmol $kg^{-1}$), (I) total alkalinity (μmol $kg^{-1}$), (J) aragonite saturation ($\Omega_{Aragonite}$). *** refers to a p-value < 0.001. ** refers to a p-value < 0.01. NS refers to not significant. See also Table S1
